# The Actin Cytoskeleton and Caveolae Regulate MERTK Cleavage by ADAM17

**DOI:** 10.64898/2026.09.29.755345

**Authors:** Rachel A Nicholson, Angela Vrieze, Bryan Heit

## Abstract

Cell surface receptors are regulated via a variety of mechanisms including proteolytic inactivation by metalloproteases. Cells often co-express both the receptor and its cognate protease simultaneously on the cell surface, often with the proteases in an active state. Despite this, receptor cleavage is generally not constitutive, and rather must be induced. How a receptor and its inactivating protease can both be present in their active form on the cell surface with no cleavage of the target receptor is unclear. Using the efferocytic receptor MERTK and its inactivating protease ADAM17 as a model, along with super-resolution microscopy approaches, we demonstrate that MERTK and ADAM17 are localized to distinct sub-regions of the plasma membrane. These microdomains are distinct subcellular structures, with MERTK contained within actin-based corrals and ADAM17 contained within caveolae. Induction of MERTK cleavage by PMA results in the reorganization of the membrane-proximal actin cytoskeleton that enables MERTK to diffuse into ADAM17-containing microdomains. Only after localizing to ADAM17-containing microdomains is MERTK cleaved. These results demonstrate that the spatial organization of the plasma membrane is dynamically regulated, with reorganization of membrane microdomains playing a role in the regulation of ADAM17 activity.

## Introduction

While often depicted as a homogeneous “sea” of lipids and proteins, the plasma membrane is instead a highly heterogeneous organelle with numerous diffusion-restricting structures enabling the segregation of proteins into distinct microdomains [1,2]. To-date, multiple membrane structures have been shown to segregate membrane proteins into distinct subregions. This includes corrals – diffusion restricting “picket-fence” structures comprised of actin “rails” and actin-bound transmembrane protein “pickets”, with the pickets acting as a percolation barrier that restricts protein diffusion out of the corral [3]. Lipid rafts can stabilize proteins into large membrane structures, for example stabilizing active T cell receptors into large multi-protein signaling synapses [4,5]. The tight association of the structural proteins that form caveolae and clathrin-coated pits with the plasma membrane sterically hinders the movement of transmembrane domains, allowing these caveolae and clathrin-coated pits to confine proteins within these structures [6,7]. Other proteins have been suggested to play a similar role in confining proteins to discrete membrane subdomains, such as tetraspanin webs [8,9]. While there are multiple structures known to restrict diffusion in the plasma membrane and to form discrete membrane microdomains, the role of these structures in regulating processes such as cellular signaling remains only partially defined.

MERTK is an efferocytic receptor that is central to the removal of apoptotic cells in many tissues [10– 13]. MERTK recognizes apoptotic cells via the opsonins Gas6, Protein S, and Tubby, which bridge MERTK to phosphatidylserine exposed on the surface of apoptotic cells [14–16]. MERTK ligation induces activation of its intrinsic kinase domain, which in-turn activates a Src-family kinase dependent signaling pathway that mediates the engulfment of the apoptotic cell [17]. MERTK utilizes integrins as co-receptors in the engulfment process, with MERTK inducing inside-out integrin activation, and the integrins providing the adhesive force required to engulf the apoptotic cell into an intracellular vacuole [17–19]. Once engulfed, the apoptotic cell is degraded and its constituent parts recycled. While over a dozen efferocytic receptors have been described, MERTK is the primary or sole efferocytic receptor in several tissues. This includes in the retina where MERTK mediates the removal of shed distal tips of photoreceptor outer segments, and in the vasculature of the heart where MERTK counteracts the accumulation of apoptotic cells into an atherosclerotic plaque [20,21]. Indeed, MERTK-inactivating mutations result in vision loss via autosomal recessive retinitis pigmentosa, and in accelerated atherosclerotic disease [22,23]. In both cases, disease is potentiated by the accumulation of apoptotic cells within the extracellular space. These uncleared cells eventually undergo secondary necrosis, spilling their inflammatory cytosolic contents into the intracellular space, leading to tissue disruption and disease progression.

In humans, MERTK function is lost in the heart independently of mutations to MERTK itself. This loss of function occurs due to the cleavage of MERTK’s extracellular domain by the membrane-bound protease ADAM17 [20,24]. Cleavage impairs efferocytosis via two mechanisms – the loss of the extracellular ligand binding domain inactivates MERTK, while the resulting soluble MERTK fragment acts as a competitive inhibitor of Gas6/MFG-E8/Tubby binding [25]. In the heart, MERTK is predominantly expressed by macrophages, which are also the primary cell type responsible for clearing apoptotic cells from this tissue [20,26,27]. These macrophages constitutively express active ADAM17 and MERTK on their cell surface, but despite this, MERTK is not constitutively cleaved by these cells and instead cleavage must be induced via inflammatory signals such as oxidized low-density lipoprotein [20]. As both MERTK and active ADAM17 are present on the cell surface at the same time, there must be additional mechanisms acting to prevent MERTK cleavage on macrophages. One potential mechanism is the segregation of MERTK and ADAM17 into distinct membrane compartments, thereby physically separating the two proteins. Our previous work has identified actin- and integrin-rich microdomains that contain MERTK, while ADAM17 has been shown to require caveolae for its activation [17,28,29]. In this study, we combine single particle tracking with super resolution microscopy to examine the segregation of MERTK and ADAM17 into discrete membrane microdomains on the cell surface. We further test the hypothesis that pro inflammatory signaling triggers MERTK cleavage by driving the coordinated mobilization of MERTK and ADAM17 out of their microdomains, thus allowing them to intermix on the cell surface.

## Results

### MERTK and ADAM17 are Non-Randomly Distributed on the Plasma Membrane

Using immunofluorescent microscopy, we investigated the distribution of MERTK and ADAM17 on the cell surface of RAW264.7 macrophages. Both MERTK and ADAM17 were present on the surface of unstimulated macrophages (**Figure 1A**), and interestingly, while both MERTK and ADAM17 coexist on the cell surface, they appear to be in spatially distinct regions of the plasma membrane. This lack of colocalization was reproduced over multiple cells (**Figure 1B**), suggesting that MERTK and ADAM17 are in spatially distinct regions of the plasma membrane. This finding raised the question of whether this spatial separation would prevent ADAM17 from cleaving MERTK in these cells. Using flow cytometry, we confirmed that 1 hour of stimulation with 50 nM PMA was able to induce ADAM17-dependent cleavage of MERTK in a PKC-dependent manner in macrophages, with the ADAM17 inhibitor KP-457 abrogating PMA-induced MERTK cleavage (**Figure 1C, S1**) [1]. To better resolve the spatial organization of MERTK and ADAM17, Ground State Depletion Microscopy (GSDM) was used to visualize these proteins with ∼20 nm resolution, confirming that these proteins are found in spatially distinct regions of the plasma membrane (**Figure 1D**). Consistent with our previous work, MERTK was found in clusters 270 ± 49 nm in diameter, while ADAM17 was found in smaller complexes of ∼160 nm ± 78 nm, with 1 hour of PMA stimulation having no effect on cluster size (**Figure 1E**) [2,3]. ADAM17 and MERTK clusters were separated by more than 900 nm – far larger than the cluster size of these proteins – confirming the spatial separation observed in conventional microscopy (**Figure 1F**), although PMA stimulation reduced the distance between clusters. Thus, while both ADAM17 and MERTK are present on the cell surface, they are localized to separate sub-regions of the plasma membrane.

**Figure 1:**
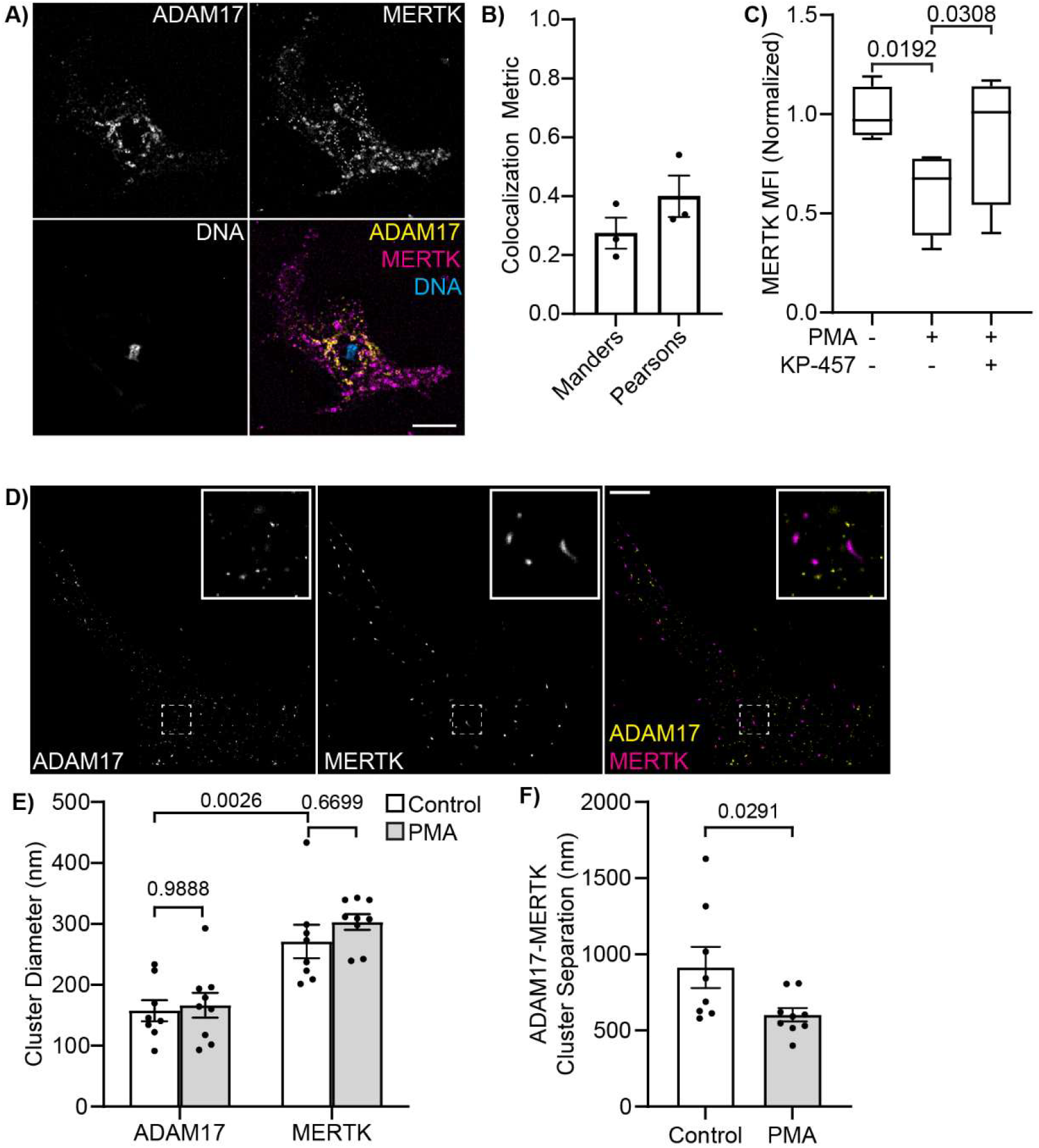
MERTK and ADAM17 Are Located on Spatially Distinct Regions of the Plasma Membrane. **A)** Fluorescence micrograph of the baso-lateral surface of a macrophage, surface-labeled for ADAM17 (yellow) and MERTK (magenta), and imaged at high magnification. Scale bar is 10 μM, DNA is stained with Hoesch (cyan). **B)** Manders colocalization coefficient and Pearson correlation coefficient between MERTK and ADAM17, quantified from fluorescent micrographs of macrophages immunostained for ADAM17 and MERTK. **C)** Flow cytometry quantification of MERTK cleavage following 1 hour stimulation with 50 nM PMA ± the ADAM17 inhibitor KP-457. **D)** GSDM image of cell surface ADAM17 (yellow) and MERTK (magenta) on the baso-lateral surface of a macrophage. Scale bar is 2.5 μM, inset shows an area 2 μm × 2 μm. MERTK and ADAM17 are localized with a precision of 20.8 ± 2.9 nm. **E-F)** ADAM17 and MERTK cluster diameter (E) and separation distance between clusters of MERTK and the nearest cluster of ADAM17 (F), quantified by GSDM imaging. n = minimum of 3 biological replicates, p values were calculated using an ANOVA with Tukey correction (C, E) or with a Welshes test (F).

### MERTK and ADAM17 Display Distinct Patterns of Diffusion

We next used Single Particle Tracking (SPT) microscopy to track the diffusion of MERTK and ADAM17 on the surface of resting macrophages. This entailed labeling MERTK and ADAM17 at an intermediate density with primary antibody, followed by fluorescently labeled secondary Fab fragments—an approach we have previously shown does not result in crosslinking of the labeled proteins [4]. After labeling, videos were recorded at a frame rate of 50 ms/frame for 300 frames, molecular trajectories were reconstructed using a multiparticle tracking algorithm, and moment scaling spectrum analysis used to identify both the mode of diffusion and diffusion coefficient [5]. Most of the MERTK and ADAM17 were found to be confined to small regions of the plasma membrane (**Figure 2A-C**). Both confined MERTK and confined ADAM17 diffused more slowly than their freely diffusing counterparts, but interestingly, freely diffusing MERTK diffused more rapidly than freely diffusing ADAM17, suggesting that ADAM17 and MERTK occupy different diffusional environments (**Figure 2D**). The confinement structures containing MERTK and ADAM17 were of different sizes, with MERTK confined to structures averaging 30 nm larger than those confining ADAM17, consistent with the observation that these proteins are localized to distinct regions of the plasma membrane (**Figure 2E**).

**Figure 2:**
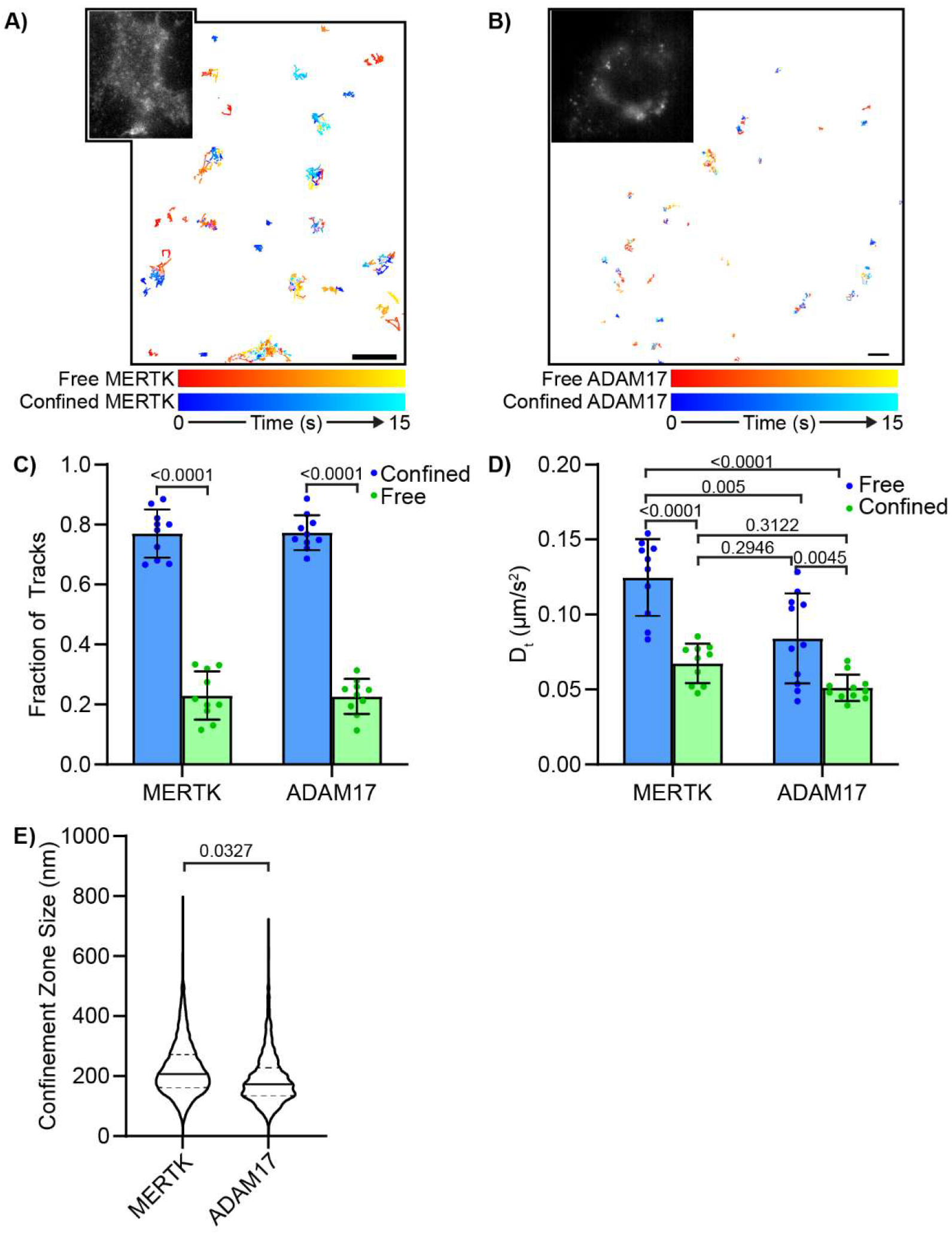
Diffusion Characteristics of MERTK and ADAM17 on Macrophages. **A-B)** Single particle tracks of MERTK (A) and ADAM17 (B) on resting macrophages, with MERTK or ADAM17 labeled at single particle tracking density using an Alexa Fluor 647 secondary antibody, and 300-frame aquations of the cells acquired at 50 ms/frame. The tracks of freely diffusing molecules are displayed using red-yellow time coding, and particles undergoing confined diffusion are displayed using blue-cyan time coding. Scale bars are 2.5 μm. Inset shows the first frame of the unprocessed immunofluorescent timelapses. **C)** Fraction of MERTK tracks undergoing confined (blue) versus free (green) diffusion. **D-E)** Diffusion coefficients (D) and confinement zone size (E) for freely diffusing (blue) and confined (green) MERTK and ADAM17. n = 10 biological repeats, p values are calculated between indicated bars using an ANOVA with Tukey correction (C, D) or Welshes test (E).

### MERTK and ADAM17 Are Confined by Distinct Membrane Structures

There are three well-described structures which have been shown to confine diffusing proteins on the plasma membrane: actin-stabilized corrals, cholesterol-stabilized lipid microdomains, and caveolae [6–8]. Corrals were disrupted by pre-treating cells with 2 μM latrunculin A – the highest concentration we could use without detaching the cells from the coverslip, while cholesterol was depleted using MβCD [9]. In untreated cells approximately 80% of MERTK and ADAM17 underwent confined diffusion, with latrunculin B treatment reducing the confinement of MERTK, and MβCD reducing the confinement of ADAM17 (**Figure 3A-B**). Moreover, the size of the confinement regions for MERTK increased following latrunculin B treatment, while the size of the confinement regions for ADAM17 increased after MβCD treatment (**Figure 3C-D**). Critically, these treatments did not result in statistically significant changes in the diffusion rate of MERTK or ADAM17, demonstrating that these changes in confinement were due to disruption of the membrane structures confining these proteins, rather than bulk changes to the diffusive environment of the plasma membrane (**Figure 3E-F**).

**Figure 3:**
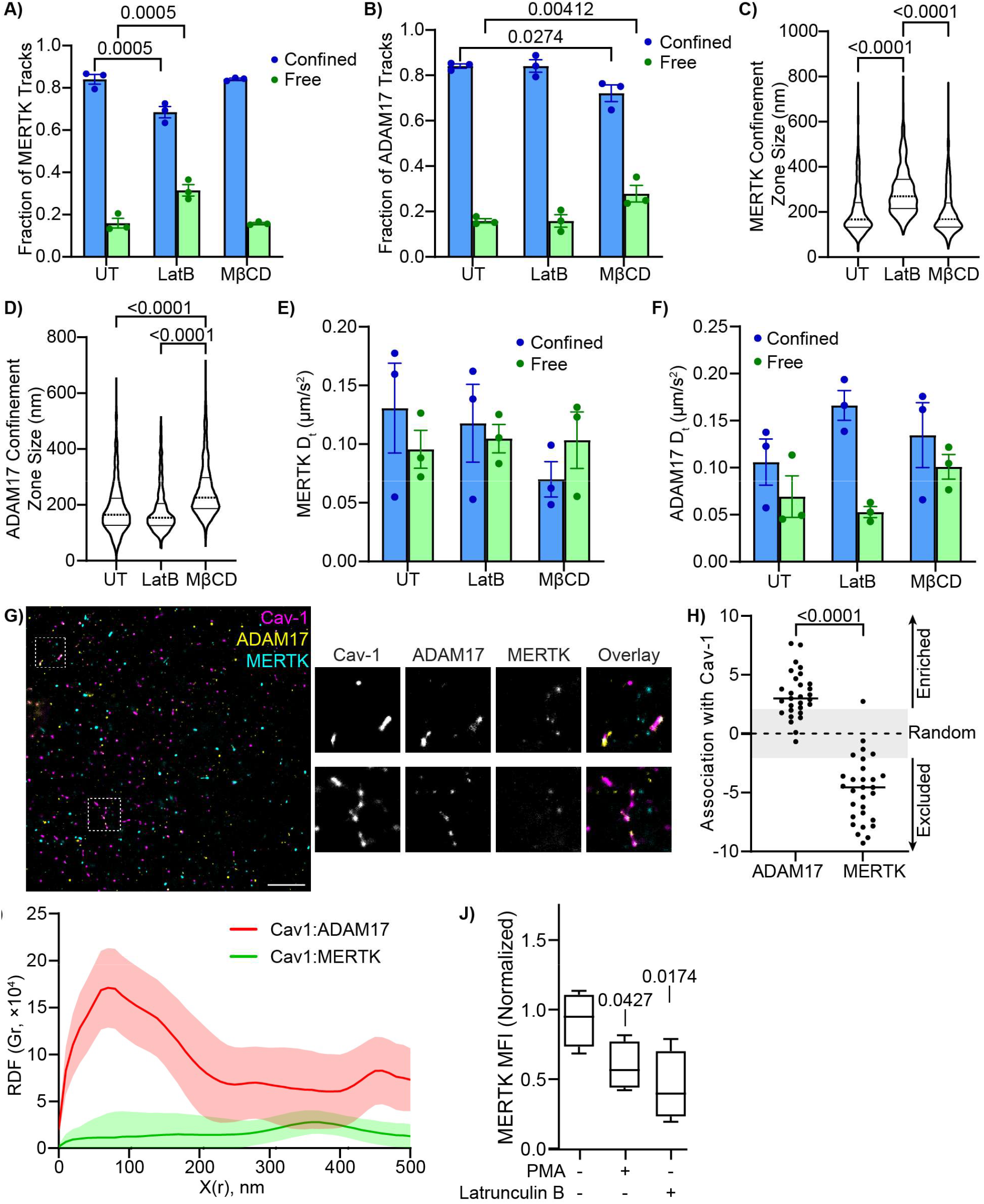
Association of MERTK and ADAM17 with Actin and Caveolae. **A-F:** Single Particle Tracking Microscopy was used to track quantify the diffusion of MERTK and ADAM17 in cells treated with Latrunculin B (LatB) to disrupt the actin cytoskeleton, methyl-β-cyclodextrin to deplete the membrane of cholesterol, or vehicle control (UT). **A-B)** Fraction of MERTK (A) and ADAM17 (B) undergoing free versus confined diffusion, **C-D)** Confinement zone size of the fraction of MERTK (C) and ADAM17 (D) undergoing confined diffusion, and **E-F)** Diffusion coefficient (D_t_) of MERTK (E) and ADAM17 (F). **G)** Ground State Depletion Microscopy was used to map the location of Cav1 (magenta), ADAM17 (yellow) and MERTK (cyan) on the surface of a macrophage. Insets show two regions where ADAM17 and Cav1 are co-clustered. Scale bar is 200 nm, localization precision is 15.4 ± 2.7 nm. **H)** Spatial Association Analysis of the nearest neighbour distances between Cav1 and ADAM17 or MERTK reveals that ADAM17 is localized closer to Cav1, and MERTK is localized further away from Cav1, than would be predicted of randomly distributed molecules. **I)** Radial Distribution Function analysis of Cav1 co-clustering with ADAM17 (Red) and MERTK (Green). **J)** Flow cytometry quantification of MERTK cleavage following stimulation with PMA or with the actin depolymerizing agent Latrunculin B. n = 30 cells imaged over 3 independent experiments (A-F), or 5 biological repeats (G-J). Data is plotted as average values of individual repeats (A,B,E,F), or ensemble data of all 30 cells (C, D, H, I). P-values are shown for statistically significant differences, 2-way ANOVA with Tukey’s test (A-F) or Mann-Whitney U test (H).

ADAM17 has previously been reported to localize to caveolae, which are also required for the activation of ADAM17 in response to some extracellular signals [7,10,11]. Caveolae have also been reported to be stabilized by cholesterol, suggesting that ADAM17 may be confined within caveolae [12]. To test this possibility, we utilized GSDM to visualize Cav1, ADAM17, and MERTK with ∼20 nm precision (**Figure 3G**). The association between MERTK and ADAM17 with Cav1 was quantified using spatial association analysis [4], demonstrating that ADAM17 was found within 20 nm of Cav1 more frequently than would be expected of randomly distributed proteins, while MERTK was excluded from Cav1 containing regions (**Figure 3H**). Consistent with cthis observation, radial distribution analysis determined that Cav1 and ADAM17 co-clustered in regions 50-100 nm in size—consistent with the size of caveolae—whereas MERTK did not co-cluster with Cav1 (**Figure 3I**). Interestingly, treating macrophages with Latrunculin B was as effective as inducing MERTK cleavage through stimulation with PMA, further suggesting that actin acts to segregate MERTK from ADAM17 (**Figure 3J**). Thus, MERTK and ADAM17 appear to be corralled by different membrane structures – MERTK by actin corrals, and ADAM17 by caveolae or another form of cholesterol-dependent microdomain.

### Changes in Corralling Control MERTK Cleavage

As the diffusion of MERTK is restricted by actin, we next ascertained whether MERTK-actin interactions changed upon PMA stimulation. Actin was labeled with SPY555-FastAct_X, MERTK or ADAM17 labeled with Alexa Fluor 647, and simultaneous high-speed timelapses of both fluorophores captured using a dichroic that projected each fluorophore onto different halves of a single camera. The SPY555-FastAct_X timelapse was used to assemble an eSRRF image with a resolution of ∼70 nm [13], on which the MERTK and ADAM17 diffusion tracks were overlayed (**Figure 4A, C, Supplemental Videos 1-3**). Significant colocalization was observed between actin and MERTK undergoing confined diffusion, but not between actin and freely diffusing MERTK or ADAM17—with PMA stimulation reducing the association of MERTK with actin (**Figure 4B, D**). PMA stimulation altered the cortical actin cytoskeleton, with the fine and highly branched network of actin fibers dissipating within 5 minutes of PMA stimulation, and later reorganizing into larger filaments (**Figure 4A, C, E, Supplemental Video 4**). The initial loss of fine actin filaments corresponded to a period where MERTK undergoes a transient increase its diffusion rate **(Figure 4F**), the total fraction of confined MERTK and ADAM17 decreased (**Figure 4G**), and where the size of the corrals confining MERTK – but not ADAM17 – expanded in size (**Figure 4H**). Combined, these results indicate that PMA stimulation results in an increase in MERTK diffusivity due to changes in the actin cytoskeleton.

**Figure 4:**
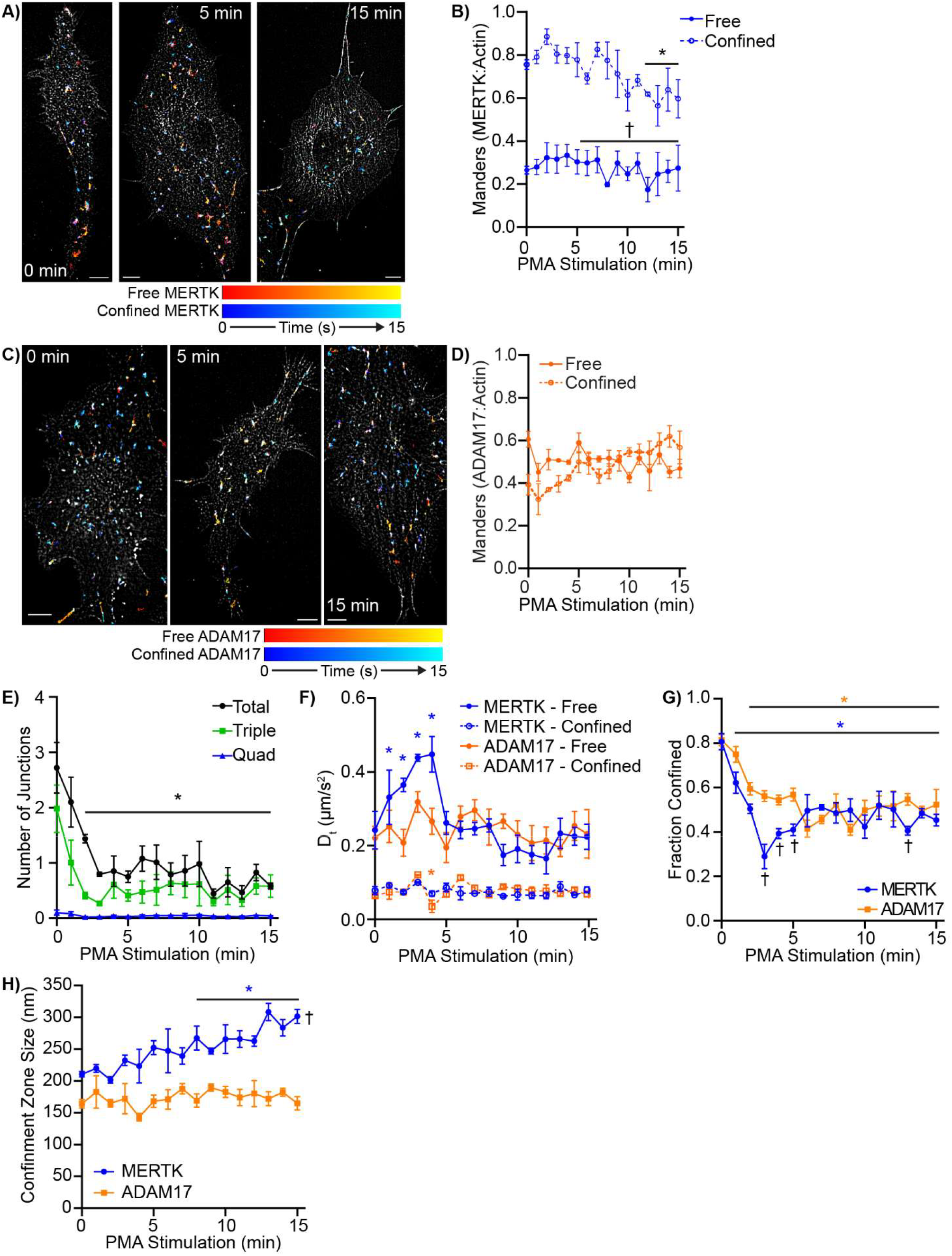
Actin Cytoskeletal Confinement of MERTK. Macrophages were labeled with the non-covalent actin dye SPY555-FastAct_X, either MERTK or ADAM17 labeled at single particle tracking density using an Alexa Fluor 647 secondary antibody, and combined SRRF-SPT imaging performed after stimulating the cells with PMA. **A)** MERTK tracks overlayed over the cortical actin cytoskeleton in cells treated for 0, 5, or 15 minutes with PMA. The tracks of freely diffusing MERTK are displayed using red-yellow time coding, and confined diffusing particles are displayed using blue-cyan time coding. **B)** Manders colocalization coefficient between actin and freely diffusing (Free) versus confined MERTK. **C)** ADAM17 tracks overlayed over the cortical actin cytoskeleton in cells treated for 0, 5, or 15 minutes with PMA. The tracks of freely diffusing ADAM17 are displayed using red-yellow time coding, and confined diffusing particles are displayed using blue-cyan time coding. **D)** Manders colocalization coefficient between actin and freely diffusing (Free) versus confined ADAM17. **E)** Quantification of actin branching following the treatment of cells with PMA. Branching is quantified as the average number of junctions per actin fiber, with total junctions, and junctions where three (triple) or four (quad) actin fibers join, quantified. PMA was added at time = 0 min. **F)** Diffusion rates of MERTK and ADAM17 following PMA treatment. PMA was added at time = 0 min. **G-H)** The fraction of MERTK and ADAM17 undergoing confined diffusion (G) and the size of the confinement zones (H) in cells following PMA treatment. n = 3 biological replicates. * p < 0.05 compared to t = 0, † p < 0.05 between MERTK and ADAM17 at the indicated timepoints (B,G) or across the entire time series (H), Kruskal-Wallis test with Dunn correction. Scale bars are 5 μm. SRRF images have a resolution of ∼75 nm, particle tracks have a localization precision better than 25 nm.

We next focused on diffusional changes 10 minutes post-PMA stimulation, as the cytoskeletal changes observed in Figure 4 were largely completed at this time. Diffusion rates of MERTK and ADAM17 had stabilized to rates similar to those observed on unstimulated cells (**Figure 5A**), as had the portion of freely diffusing versus confined diffusing MERTK and ADAM17 (**Figure 5B**). As observed previously, ADAM17 was confined in regions smaller than those confining MERTK (**Figure 5C**). Unexpectedly, there was a slight but statistically significant decrease in the size of MERTK confinement zones following PMA stimulation. This did not appear to be a monotonic downward shift in the size of the MERTK confinement zones, and rather a second smaller confinement zone population seems to have appeared (**Figure 5C**). To determine if a second population had emerged, we used a Gaussian Mixture Models approach to attempt to separate these populations. While single log-normalized Gaussian best modeled the confinement zone sizes of ADAM17 and of MERTK in untreated cells, a two Gaussian model best reproduced the distribution of confinement zone sizes observed in the PMA-treated MERTK group (**Figure 5D**). Interestingly, the first Gaussian had a similar mean and distribution to MERTK in untreated cells, while the mean and distribution of the smaller Gaussian resembled that of confined ADAM17. This suggests that a portion of MERTK may be entering the confinement zones that normally contain ADAM17.

**Figure 5:**
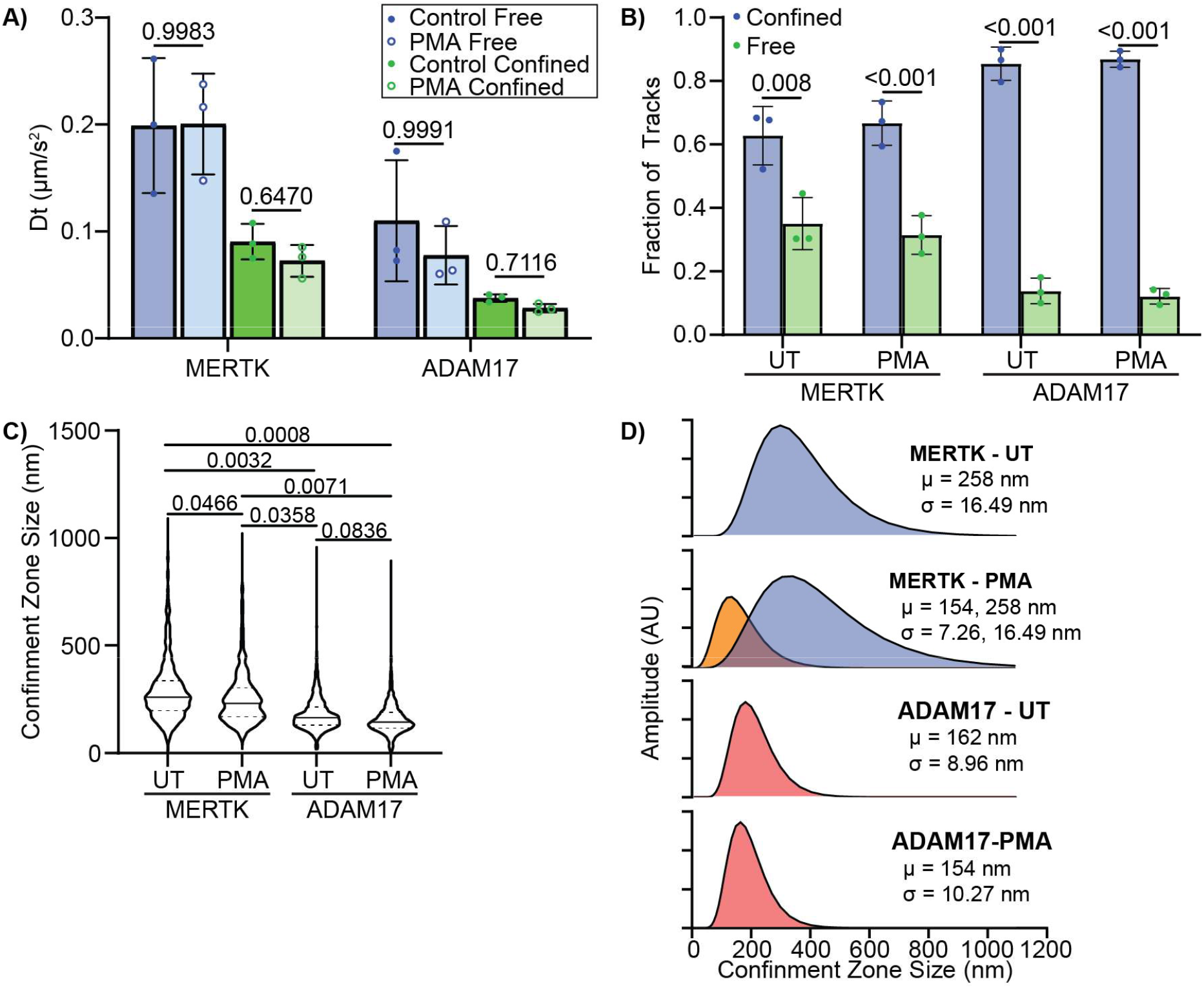
MERTK Confinement Changes Following PMA Stimulation. Macrophages were stimulated with 50 nM PMA for 10 minutes, then single particle tracking was used to characterize differences in MERTK and ADAM17 diffusion. **A-B)** Changes in the diffusion rate (A) and fraction of molecules undergoing free versus confined diffusion (B) in unstimulated (UT) versus PMA-stimulated (PMA) macrophages. **C)** Distribution of MERTK and ADAM17 confinement zone sizes in unstimulated (UT) versus PMA-stimulated (PMA) macrophages. **D)** Gaussian mixture model decomposition of the MERTK and ADAM17 distributions from panel C. The distribution of confinement zone size for all treatments other than MERTK in PMA-treated cells (MERTK-PMA) could be described as a single log-normalized Gaussian, whereas the distribution of MERTK-PMA was best described by the sum of two Gaussians. The mean (μ) and standard deviation (σ) for each distribution are shown. n = 3 (A-C) or composite data from all three experiments (D). p-values were calculated using a 2-way ANOVA with Tukey’s test.

To investigate whether cleavage occurred within ADAM17 confinement zones, we used the “split camera” approach from above to simultaneously track MERTK and ADAM17 on the surface of PMA-stimulated macrophages, defining putative MERTK-ADAM17 interactions as any time the inter-molecular distance dropped below our precision of detection (25 nm) for more than 1 frame. As expected, on unstimulated cells MERTK and ADAM17 were found to be highly confined in neighbouring – but not intermixing – corrals (**Figure 6A)**. Occasional interactions between MERTK and ADAM17 were observed (**Figure 6A**, inset), but these tended to be transitory interactions of short duration (**Figure 6B**). We defined putative cleavage events as those where a MERTK track ends following an interaction with ADAM17, prior to the end of the ADAM17 track. On resting macrophages, less than 1% of MERTK exhibited putative cleavage events (**Figure 6E**). PMA stimulation increased the frequency of MERTK-ADAM17 interactions on macrophages (**Figure 6C**). While many of these interactions were transitory, there were a marked increase in the number of interactions which led to putative MERTK cleavage events (**Figure 6C, E**). In both untreated and PMA-stimulated macrophages, MERTK-ADAM17 interactions that led to putative cleavage events were longer in duration than those which did not result in cleavage (**Figure 6F**). Interestingly, cleavage always involved a MERTK that was co-confined with ADAM17 at the start of the observation period, or a MERTK that transitioned from freely diffusing to co-confined with ADAM17 prior to cleavage (**Figure 6C** inset, **6F**). These data indicate that MERTK is cleaved by confined ADAM17, that cleavage requires a modest (∼0.5 second) interaction time between MERTK and ADAM17, with PMA stimulation increasing the frequency of MERTK-ADAM17 intermixing and cleavage through mobilizing MERTK out of actin-based corrals and into caveolae-sized structures containing ADAM17.

**Figure 6:**
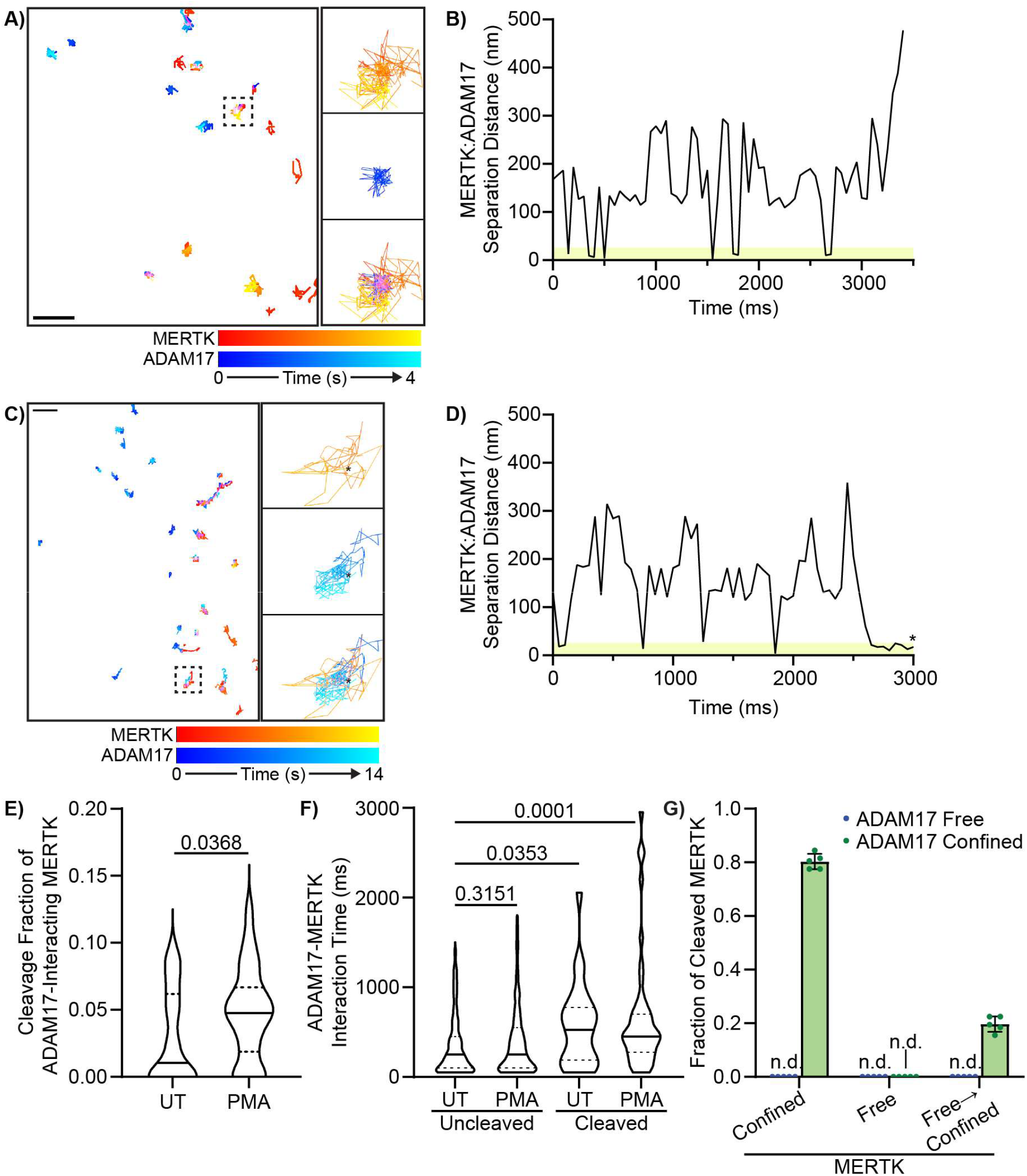
MERTK Cleavage is Mediated by Confined ADAM17. Dual-colour single particle tracking was used to quantify interactions between MERTK and ADAM17 diffusing on the cell surface. MERTK-ADAM17 interactions were defined as events where MERTK and ADAM17 were detected 25 nm of each other, with MERTK, with putative MERTK cleavage events defined as the loss of a MERTK following an interaction with ADAM17. **A)** Representative diffusion tracks of MERTK (red-yellow) and ADAM17 (blue-cyan) on a resting macrophage. Inset show a single MERTK and ADAM17 track that are diffusing in the same general region of the cell. **B)** MERTK-ADAM17 separation distance from the molecules tracked in the inset from panel A. Putative interactions occur when the distance between molecules is less than 25 nm (shaded region). **C)** Representative diffusion tracks of MERTK (red-yellow) and ADAM17 (blue-cyan) on a PMA-stimulated macrophage. Inset show a single MERTK and ADAM17 track that are diffusing in the same general region of the cell. **D)** Quantification of MERTK:ADAM17 separation distance from the molecules tracked in the inset from panel C, displaying a putative cleavage event (*). **E)** Frequency of non-cleavage versus putative cleavage events following MERTK-ADAM17 interactions on untreated versus PMA-simulated cells. **F)** Interaction times for non-cleavage versus putative ADAM17-MERTK cleavage events on untreated versus PMA-simulated cells. **G)** Fraction of cleaved MERTK that was diffusing in a Confined, Free, or became confined after freely diffusing (Free⟶Confined) by ADAM17 which was freely diffusing versus undergoing confined diffusion. n.d. = none detected. Data is plotted as a single representative track (B,D), or as mean +/-SEM (E-G), p-values were calculated using a Mann-Whitney u test (E) or ANOVA with Tukey correction (F-G). Scale bars are 0.5 μm.

## Discussion

MERTK is the predominant efferocytic receptor expressed in many tissues, where it mediates the recognition and engulfment of apoptotic cells [14– 17]. Under some inflammatory conditions, including atherosclerosis and ischemia-reperfusion injury, the extracellular domain of MERTK is cleaved by the protease ADAM17, releasing a soluble fragment [18,19]. This has a profound inhibitory effect on efferocytosis in the inflamed tissue, as cleavage inactivates MERTK, while the soluble fragment acts as a competitive inhibitor of the efferocytic function of not only MERTK, but also of the related efferocytic receptors AXL and TYRO3 [20]. This loss of efferocytic activity enhances disease development and tissue injury in these patients, worsening outcomes. While the role of MERTK and efferocytosis in these conditions are well known, the mechanisms regulating MERTK cleavage by ADAM17 remain only partially understood. In this study we have demonstrated that MERTK and ADAM17 are segregated into distinct regions of the plasma membrane, with MERTK confined to actin-rich corrals, and ADAM17 confined within caveolae. Stimulation of these cells with PMA induced the ADAM17-dependent cleavage of MERTK, with cleavage enabled by cytoskeletal rearrangements that release MERTK from corrals, allowing it to move to ADAM17-containing microdomains where it is cleaved. These data demonstrate the important role of membrane microdomains in regulating protease-ligand interactions, and demonstrate that the spatial structuring of proteases and their ligands on the cell surface is part of the cellular machinery that regulates proteolysis.

Confinement within actin-based corrals is not unique to MERTK; several phagocytic receptors including CD36, FcγRI, FcγRIIA, FcεRI, α_m_β_2_, and RAGE have likewise been shown to reside in corrals or related actin-dependent diffusive barriers [21–26]. For these receptors, corrals appear to serve several purposes. Treatment of cells with actin-disrupting compounds such as Latrunculin B results in the ligand-independent clustering of some receptors, suggesting that corrals may act to limit spontaneous receptor aggregation [27]. Conversely, corrals also mediate for the formation of receptor microclusters – a phenomenon which increases receptor avidity and activity, indicating that corrals “fine tune” the size of receptor clusters [28–30]. The tuning of receptor clustering also occurs during active signalling, for example, in phagocytosis corrals trap Fcγ receptors along the leading edge of the phagocytic cup which both enhances the ligation of FcγR and facilitates phagosome formation [24]. In addition to structuring receptors, corrals also control spatial relationships between receptors, co-receptors, and inhibitory receptors. For example, we recently demonstrated that corrals pre-structure MERTK into microdomains containing its co-receptor α_x_β_2_ and several signaling molecules, while other studies have shown that co-clustering of receptors with the inhibitory phosphatase CD45 fine-tunes their sensitivity to ligation by raising the “noise floor” that must be overcome before productive receptor signaling occurs [2,31]. Our results expand on this regulatory role, demonstrating that corrals can regulate receptor function by controlling a receptors susceptibility to proteolysis. Beyond regulating receptor activity, corrals also participate directly in force generation during endocytosis, where the meshwork of corrals provides a compressive scaffold that helps overcome cellular turgor pressure to accelerate membrane invagination [32,33].

ADAM17—also known as Tumour Necrosis Factor-α Converting Enzyme (TACE)—is synthesized in the ER as an inactive zymogen whose pro-domain keeps ADAM17 inactive during synthesis. After synthesis, ADAM17 associates with its obligate binding partners— iRhom1 and iRhom2—which regulate ADAM17 trafficking to the cell surface, and which enable the removal of the pro-domain by furin in the trans-Golgi network [34–36]. Once at the cell surface, the catalytic activity of ADAM17 is dynamically regulated though mechanisms still under investigation. Le Gall *et al*. found that the activation of ADAM17 activity was independent of its cytoplasmic domain, and can occur without removal of its pro-domain [37]. While ADAM17 activity can be upregulated via exocytosis of intracellular ADAM17 stores, this is not required, with G-protein coupled receptor signaling inducing ADAM17 activity without changes in cell-surface ADAM17 levels [38]. ADAM17 contains a phosphatidylserine binding motif within its membrane-proximal extracellular domain, with the flipping of phosphatidylserine from the cytosolic to exofacial side of the plasma membrane activating ADAM17 [39,40]. While phosphatidylserine flipping is most often associated with apoptosis, it can also occur in response to a variety of stimuli independently of cell death [41–44]. PMA increases cell surface phosphatidylserine levels directly through PCK-mediated activation of scramblase, and indirectly by inducing the endocytosis of the flippase ATP11c [45,46]. While phosphatidylserine is the best understood activator of ADAM17, additional processes control the activity of this protease. Two recent studies have found that iRhom1 and iRhom2 remain closely associated with ADAM17 on the cell surface, where movement of the re-entry loop—a cytoplasmic amphipathic α-helix found in both iRhom proteins—acts as a molecular relay that activates ADAM17 in response to intracellular signals [47,48]. Whether re-entry loop-mediated activation is independent of phosphatidylserine is currently unknown.

ADAM17 activation is also location-dependent, with ADAM17 co-precipitating with the primary structural protein of caveolae, caveolin-1 [49]. This association with caveolae appears to be required for ADAM17 activity. Moreno-Càceres *et al* found that caveolae were required for the TGFβ-mediated induction of ADAM17 activity via supporting a Src-dependent signaling pathway [50]. In adipocytes, destabilization of caveolae by knockdown of Cavin-3 impaired the ADAM17-dependent shedding of preadipocyte factor-1 (TNFR1), while caveolin-1 knockdown in vascular endothelial cells prevents ADAM17-mediated sheading of tumour necrosis factor receptor 1 [10,11]. Our results are consistent with these findings: we found ADAM17 localized to caveolae in resting cells, with PMA stimulation resulting in no apparent change to this distribution despite a measurable increase in ADAM17 activity. This suggests that ADAM17 activation and activity occurs within caveolae, and that increases in the mobility of its target proteins are what provides ADAM17 access to its substrates. A similar spatial mechanism has been described for the chemokine CX3CL1, whose cleavage by ADAM10 is constrained by the confinement of CX3CL1 within actin corrals—although the subcellular localization of ADAM10 remains unclear [51]. Corrals are not the only cellular structure that restricts ligand interactions with ADAM-family proteases – membrane cages, a lipid-based domain distinct from rafts, confine the diffusion of CD93 on resting cells, with inflammatory stimuli inducing ADAM-dependent cleavage to form the opsonin sCD93 [9,52]. In addition to caveolae, two other mechanisms may enable localized activation of ADAM17 in discrete regions of the plasma membrane. While processes such as apoptosis and PMA-induced PKC activity result in the exposure of phosphatidylserine across the entirety of the plasma membrane, several receptors have been identified that induce highly localized phosphatidylserine flipping [53]. This could potentially induce ADAM17 activity that is localized to regions proximal to active receptors, allowing ADAM17 to selectively cleave these receptors—and indeed, microdomain-localized ADAM17 activity has been shown to mediate histamine-induced TNFR1 sheading on vascular endothelial cells [10]. In addition, the focal exocytosis which occurs during phagocytosis may deliver intracellular stores of ADAM17 directly to forming phagocytic cups, potentially providing selectivity to the inhibitory role of ADAM17 on phagocytosis [54,55].

While our study provides evidence that changes in the diffusional environment on the cell surface regulates ADAM17 activity, there are several limitations to this work. While PMA is a potent inducer of ADAM17 activation, it does so throughout the entirety of the cell membrane and affects the entirety of the cell cytoskeleton. While this type of activation is observed in response to some ADAM17-inducing conditions such as apoptosis, other ADAM17-inducing stimuli are known to produce focal externalization of phosphatidylserine and lesser impacts on the actin cytoskeleton, and therefore may drive more localized activation of ADAM17 [53,56]. In addition, ADAM17 activity can also be increased by the exocytosis of preformed stores, and as this ADAM17 would not be labeled, we would not have captured any influence of exocytosed ADAM17 in our particle tracking experiments [38]. Actin corrals undergo transient disassembly at sites of exocytosis, and as such, some of the changes in MERTK diffusivity that we observed may be due to exocytosis-mediated, rather than PMA-mediated, cytoskeletal changes [57]. Lastly, we cannot separate bona fide cleavage events from photobleaching events in our single particle tracking assays, although the use of stable fluorophores and optimization of our imaging conditions to produce <1% photobleaching should minimize this as a confounding factor [58,59].

In conclusion. this study has identified a mechanism by which cells use membrane microdomains to segregate proteases and their ligands to limit ligand cleavage, with release of ligands from these microdomains – but not release of the protease – then enabling ligand-protease interactions and ligand cleavage. Although our study focused on ADAM17 and its substrate MERTK, prior work has shown that ADAM17 similarly cleaves TNFR1 within membrane microdomains, and that spatial confinement of ligands in actin corrals limit ADAM10’s proteolytic activity [10,51]. These and our studies indicate that membrane organization governs the accessibility and activity of membrane-bound proteases, and that microdomain reorganization is part of the regulatory regime that controls proteolysis on the cell surface.

## Materials & Methods

### Materials

THP-1 and RAW264.7 cells were obtained from Cedar Lane Labs (Mississauga, Canada). All antibodies used in this study are described in **Supplemental Table 1**. Roswell Park Memorial Institute (RPMI), Dulbecco’s modified Eagle’s medium (DMEM) and fetal bovine serum (FBS) were purchased from Wisent (Saint-Jean-Baptiste, Canada), trypsin-EDTA and antibiotic-antimycotic were purchased from Corning (Manassas, Virginia, USA). KP-457, Gö6983, Doramapimod and Latrunculin B were purchased from Cayman Chemical (Ann Arbour, USA). Human TruStain FcX was from Biolegend (San Diego, USA). Round coverslips (#1.5 thickness 18 mm diameter) and 16% paraformaldehyde (PFA) were purchased from Electron Microscopy Supplies (Hatfield, Pennsylvania, USA). All tissue culture plates, centrifuge tubes, plastic ware, Hoechst 33342, and Permafluor mounting medium were purchased from Thermo Fisher Scientific. SMART dSTORM buffer was purchased from Abbelight (Cachan, France). SPY555-FastAct was from Spirochrome (Stein am Rhein, Switzerland). Flexible UV resin and UV torch were purchased from Let’s Resin (Dallas, USA). All other chemicals were purchased from Bioshop Canada (Mississauga, Ontario). Partek Flow was purchased from Parteck (St. Louis, USA), MATLAB software was purchased from MathWorks (Natick, USA), and Prism software was purchased from and GraphPad (La Jolla, California, USA). FIJI was downloaded from https://fiji.sc/ [60].

### Cell Culture

RAW264.7 macrophages were grown in DMEM + 10% FBS in a 37°C/5% CO_2_ incubator until ∼80% confluent, and then split by scraping the cells into suspension and replating at a 1:5 dilution. THP-1 monocytes were grown in RPMI + 10% FBS in a 37°C/5% CO_2_ incubator and maintained at a concentration between 500,000 – 1,000,000 cells/mL. For imaging experiments, 18 mm diameter circular coverslips, measured with a micrometer to be between 0.165 and 0.175 mm in thickness, were placed into the wells of a 12-well plate and 250,000 cells/well added to each well. Cells were the cultured at 37°C/5% CO_2_ for 24 hours prior to imaging. For flow cytometry, 500,000 THP-1 cells per well were seeded into a 12 well plate in RPMI + 10% FBS+ 100 nm of PMA and incubated at 37°C/5% CO_2_ for 48 hours. After 48 hours the PMA containing media was replaced with fresh RPMI + 10% FBS.

### Fluorescence Microscopy

RAW264.7 or THP-1 macrophages were cultured on coverslips as described above and were fixed for 15 min at 37°C in PEM buffer (80 mM PIPES, 5 mM EGTA, 2 mM MgCl2, pH ∼6.9) plus 4% PFA to limit fixation-induced receptor clustering [61]. For surface staining, cells were blocked for 30 min with PBS + 0.5% casein, and then labeled at saturation with anti-MERTK and anti-ADAM17 in PBS + 0.35% casein (**Supplemental Table S1**). The cells were then washed 3 × 5 min in PBS and labeled with Cy3- and Alexa Fluor-647-labeled secondary Fab fragments, which we have shown previously does not induce antibody-mediated receptor clustering [4]. The cells were then washed 3 × 5 min in PBS, counterstained with Hoechst 33342 for 5 min, and then mounted on a slide using Permafluor mounting media. The cells were mounted with Permafluor and imaged using a Zeiss AxioObserver equipped with a 100×/1.40 NA oil objective, Hamamatsu W-View Gemini image splitter, coupled to a Hamamatsu ORCA-Fusion CMOS camera, operated using Zeiss Zen Blue software. Full z-stacks of cells were captured progressing from longest to shortest wavelength with a 0.24 μm step size. To reduce image blur and improve the precision of colocalization analyses, the Zeiss Deconvolution toolkit was applied utilizing a constrained iterative method. MERTK and ADAM17 colocalization and cross-correlation was quantified in FIJI using Just Another Colocalization Plugin [62].

### Single Particle Tracking Microscopy

Cells cultured on coverslips were cooled to 10°C to stop endocytosis without inducing actin depolymerization [63]. Culture media was then exchanged for imaging buffer (150 mM NaCl, 5 mM KCl, 1 mM MgCl_2_, 100 µM EGTA, 2 mM CaCl_2_, 20 mM HEPES). Cells were labeled at non-saturating concentrations with anti-MERTK and/or anti-ADAM17 in imaging buffer + 1% BSA for 10 min at 10°C. The cells were washed 3 times with imaging buffer, labeled with ATTO550- and/or Alexa Fluor 647 secondary Fab fragments in imaging buffer + 1% BSA for 10 min at 10°C. The cells were washed 3 times with imaging buffer and then immediately transferred to the heated/CO_2_ perfused stage of the Zeiss AxioObserver and imaged using the 100×/1.40NA objective lens. For single-colour imaging, 300 frame time-lapse images were captured using burst mode, 50 ms acquisition time, and 2× binning. These conditions were selected as they produced less than 1% photobleaching in control experiments (not shown). For dual-colour imaging, prior to each experiment the W-View image splitter was configured to split ATTO550 and Alexa 647 fluorescence and aligned using a slide containing 200 nm diameter tetraspeck beads. Following alignment, 300 frame time-lapse images were captured using burst mode, 50 ms acquisition time, and 2× binning. The resulting dual-colour images were then denoised using Noise2Void, a pixel-level x/y alignment performed, and the aligned images exported as single-channel images using a custom macro in Zeiss Zen Blue software [64].

The positions of individual MERTK and ADAM17 molecules were then mapped with supper-resolution precision, and linked into molecular trajectories, using the approach of Jaqaman *et al*., via custom-written MATLAB scripts [5]. Molecular trajectories with a positional accuracy worse than 25 nm were filtered from the dataset, then moment scaling spectrum (MSS) analysis used to classify the diffusive patterns of individual molecules based on the power-law indices (y_ρ_) of the first through fourth moments of molecular movement (ρ), defining Free diffusion as diffusion approximating Brownian motion (ρ≈y_ρ_), and subdiffusive (confined) movement as ρ<y_ρ_ [65]. The diffusion rate is estimated from the initial slope of mean-squared displacement plot of each particle, and for confined particles, the confinement zone diameter is calculated as the 90_th_ percentile of the maximum possible extent of the particles variance-covariance matrix. Gaussian mixed models were applied in Graphapad prism to the distribution of confinement zone sizes to identify the presence of multiple confinement zone populations within single cells.

### Ground State Depletion Microscopy

Cells cultured on coverslips were fixed for 15 min at 37°C in PEM buffer + 4% PFA to limit fixation-induced receptor clustering [61]. The cells were then blocked for 10 min with 0.5% casein in PBS, then MERTK and ADAM17 labeled at saturation in PBS + 0.5% casein for 20 min (**Supplemental Table S1**). Cells were washed 3 × 5 min in PBS, then labeled with Alexa fluor 488, 555 or 647 secondary Fab fragments, for 20 min in PBS + 0.5% casein. The fluorophore used to label each protein was changed in each experimental repeat to avoid any artefacts from differences in label-specific blinking dynamics. The cells were then washed 3 × 5 min in PBS and antibody labeling stabilized by a secondary fixation using 2% PFA in PBS for 5 min. For experiments co-labeling Cav1, cells were labeled for surface ADAM17 and MERTK as above and then permeabilized with 0.1% triton X-100 in PBS + 0.5% casein for 45 min, followed by addition of anti-Cav1 in PBS + 0.5% casein for 30 min. The cells were washed 3 × 10 min in PBS, then secondary Fab added for 30 min in PBS + 0.5% casein. The cells were then washed 3 × 20 min, secondary fixation with 2% PFA in PBS for 5 min performed, and the cells washed 3 × with PBS. The coverslip was immediately transferred onto a depression slide containing Abbelight Smart Buffer and sealed to the slide using UV-curable resin. The slide was imaged on a Leica GSDM microscope, imaging the basolateral cell surface of the cell. The resulting molecular coordinate lists were imported into our MIiSR analysis software where spatial association analysis was used to quantify inter-protein interactions, and radial distribution analyses was used to quantify co-clustering [4]. To confirm that these associations were not spurious, Monte Carlo simulations were used to mimic non-interacting proteins by randomizing protein positions over the same image area, and these analyses repeated.

*Actin Imaging with Super Resolution Radial Microscopy* RAW264.7 cells were split onto coverslips as described above. SPY555-FastAct_X was diluted in 50 μL of anhydrous DMSO, as per manufactures instructions, a 1:1500 dilution added to the cells, and the cells incubated for 1 hr at 37°C and 5% CO_2_. For some experiments, the cells were then cooled to 10°C and labeled for single particle tracking of either MERTK or ADAM17 and an Alexa Fluor 647 secondary antibody, as described above. The cells were then transferred to a Leiden chamber containing imaging buffer and a 1:2500 dilution of SPY555-FastAct_X, and transferred to the heated stage of our Zeiss AxioObserver and imaged using the 100×/1.40NA objective lens. An initial acquisition was acquired, and then 50 nM of PMA added to the cells, with cells then imaged every 60 sec for 15 min. For single-colour imaging, the full sensor was acquired using 300 frame time-lapse images were captured using burst mode, 50 ms acquisition time, no binning. The resulting time series were then processed to generate enhanced SRRF images using a custom python script and the NanoPyx python package [13], with all reconstructions performed using magnification = 5, ring radius = 2.5, and sensitivity = 2.5. To quantify actin morphology, these SRRF images were segmented in Ilastik using a trained pixel classification routine, the segmented images imported into FIJI and analyzed using the Skeletonize 2D/3D and AnalyzeSkeleton plugins [60,66,67]. For combined SRRF/SPT imaging, images were acquired using the same acquisition settings as for dual-colour SPT, described above. The SRRF (SPY555 channel) and SPT (Cy5 channel) images were then split, the SRRF image reconstructed, and SPT analysis performed, as described above. The X and Y coordinates of each tracked particle was exported into .csv files, separating tracks for free versus confined tracks into different files. A custom python script used to generate time-coded tracking images with the same pixel scaling as the corresponding SRRF images (26 nm/pixel). These were then overlayed on the SRRF images and both Manders colocalization coefficients and Van Steensel’s Cross-Correlation Function analysis performed to quantify the spatial relationship between actin and confined verses freely diffusing tracks, using the Just Another Colocalization Plugin in FIJI [62].

### MERTK Shedding and Flow Cytometry

THP-1 cells were differentiated into macrophages, treated with 10 µM of KP-457, 1 µM of Doramapimod or 10 µM of Gö6983 for 30 min and then stimulated with 50 nM PMA for one 1 hr, or with 2 µM of Latrunculin B for 30 min. Cells were then washed and collected using PBS + 5mM EDTA to lift the cells. The cells were then stained with fixable viability dye eFluor450 for 30 min on ice. After washing, the cells were blocked with Human TruStain FcX blocker for 10 min at room temperature and then stained with human anti-MERTK-APC antibody for 30 min at room temperature (Supplemental Table S1). After washing, cells were fixed with 4% PFA in PBS for 15 min at room temperature. After washing, the cells were resuspended in FACS buffer (2% FBS + 5 mM EDTA in PBS) and a minimum of 10 000 cells acquired on a BD FACSymphonyA1. FlowJo was used to analyze the resulting data by first gating on forward and side scatter, selecting singlets with the forward scatter height/area channels, and live cells gated via the eF450 channel. A ZsGreen+ gate was then used to confirm the cell line. The median fluorescence intensity for MERTK was then determined for MERTK+ population.

### Data Transparency

All custom Matlab and Python scripts, as well as the trained Ilastik model for segmenting actin SRRF images, are available on the Heit Lab Github repository (https://github.com/bheit/ADAM17-MERTK). All numerical data are included in the study, and the microscopy images are available upon request to the corresponding author.

### Statistical Analysis

Unless otherwise indicated data are presented as the mean ± s.e.m. and analyzed using a two-tailed unpaired Student’s t-test or one-way ANOVA with Tukey correction. All statistical analyses were performed in GraphPad Prism Version 11.

## Acknowledgments

We would like to thank the Molecular Imaging Facility for their assistance in the single-particle tracking, SRRF, and GSDM microscopy, and to thank the London Regional Flow Cytometry Facility for their assistance with the flow cytometry experiments.

## Supplemental Materials

**Supplemental Figure 1:**
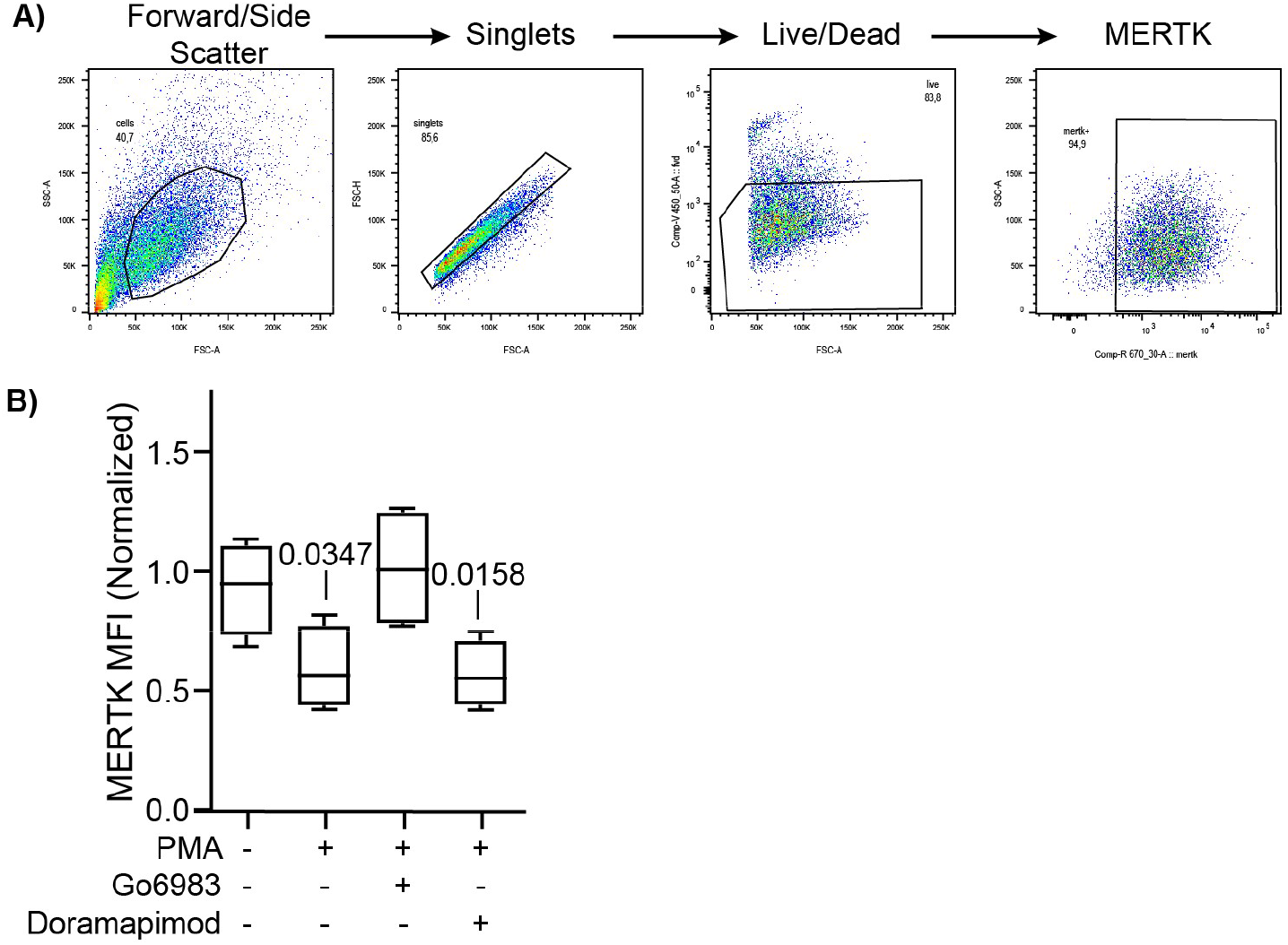
Flow Cytometry Quantification of MERTK Cleavage. **A)** Gating strategy used to quantify MERTK on macrophages. Forward/side scatter is used to identify cells matching the size and granularity of macrophages, scatter height and area used to detect singlets, living cells selected by gating for cells negative for violet viability due, and then MERTK quantified on the resulting selected cell population. **B)** Impact of PKC (Go6983) and p38 MAPK (Doramapimod) inhibitors on PMA-induced MERTK shedding. n = minimum of 3 biological replicates. P-values were calculated using ANOVA with Tukey correction.

**Supplemental Table 1:**
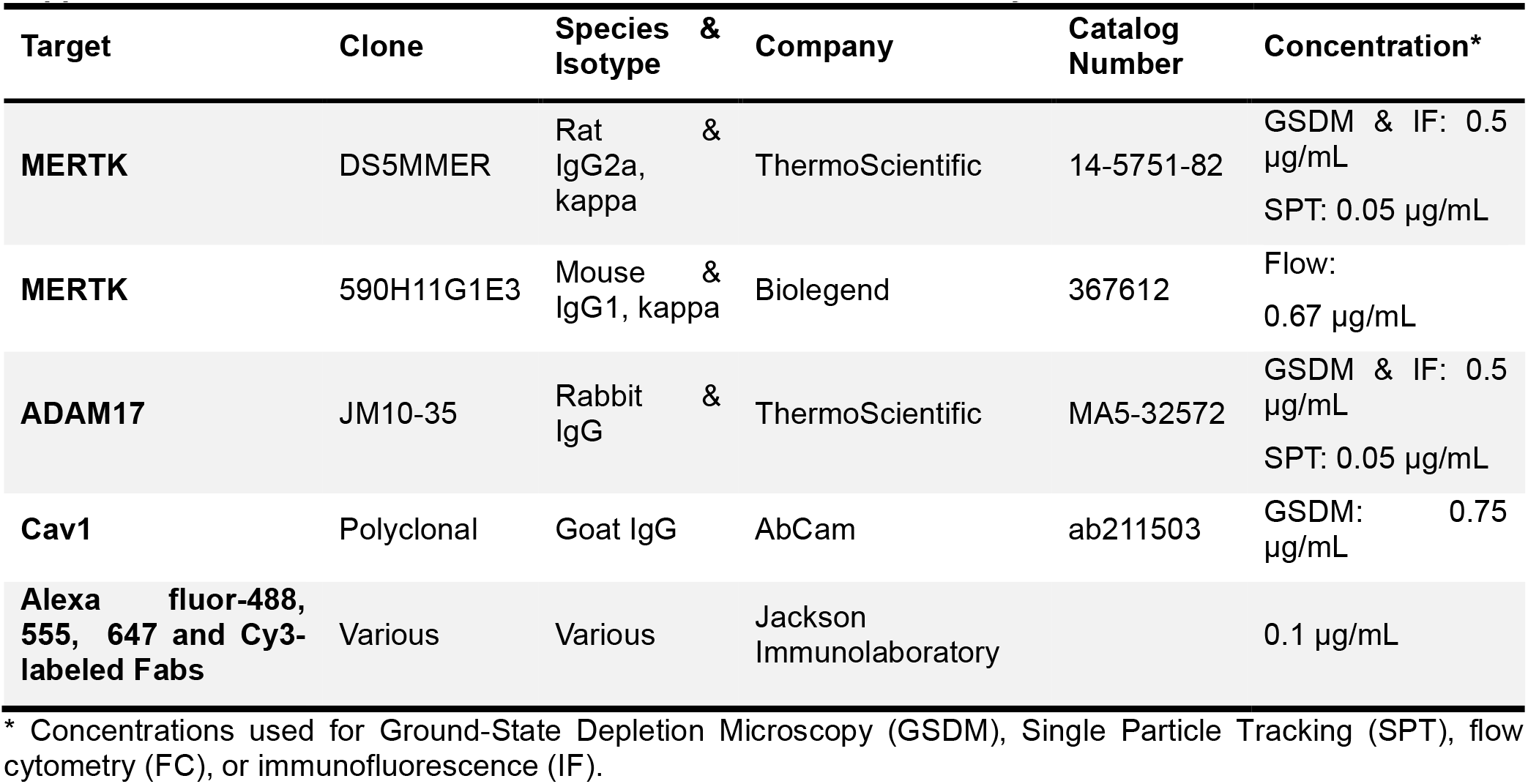
Antibodies and Other Stains Used in This Study.

**Supplemental Video 1: Overlay of MERTK and ADAM17 Diffusion Tracks on Macrophages Prior to Stimulation with PMA**. Macrophages were labeled with the non-covalent actin dye SPY555-FastAct_X, either MERTK or ADAM17 labeled at single particle tracking density using an Alexa Fluor 647 secondary antibody, and combined SRRF-SPT imaging performed. The tracks of freely diffusing MERTK are displayed using red-yellow time coding, and confined diffusing particles are displayed using blue-cyan time coding. https://youtu.be/O2V1QomK7nM

**Supplemental Video 2: Overlay of MERTK and ADAM17 Diffusion Tracks on Macrophages After 5 Minutes of PMA Stimulation**. Macrophages were labeled with the non-covalent actin dye SPY555-FastAct_X, either MERTK or ADAM17 labeled at single particle tracking density using an Alexa Fluor 647 secondary antibody, stimulated with 50 nM PMA for 5 minutes, and combined SRRF-SPT imaging performed. The tracks of freely diffusing MERTK are displayed using red-yellow time coding, and confined diffusing particles are displayed using blue-cyan time coding. https://youtu.be/FiYXfM7oX1U

**Supplemental Video 3: Overlay of MERTK and ADAM17 Diffusion Tracks on Macrophages After 15 Minutes of PMA Stimulation**. Macrophages were labeled with the non-covalent actin dye SPY555-FastAct_X, either MERTK or ADAM17 labeled at single particle tracking density using an Alexa Fluor 647 secondary antibody, stimulated with 50 nM PMA for 15 minutes, and combined SRRF-SPT imaging performed. The tracks of freely diffusing MERTK are displayed using red-yellow time coding, and confined diffusing particles are displayed using blue-cyan time coding. https://youtu.be/ozh9kgw7T8g

**Supplemental Video 4: Actin Dynamics Following PMA Stimulation**. Macrophages were labeled with the non-covalent actin dye SPY555-FastAct_X, stimulated with 50 nM PMA, and SRRF images acquired every minute for 15 minutes. The cytoskeleton reorganizes from a fine meshwork to larger fibers, and multiple actin-bound endosomes can be observed. https://youtu.be/g-0vdk6NdfM

